# Regional connectivity and viability selection in a range-expanding marine species

**DOI:** 10.64898/2026.03.28.714283

**Authors:** Andy Lee, Benjamin N. Daniels, Cataixa López, Jean M. Davidson, Robert J. Toonen, Mark R. Christie, Crow White

## Abstract

Identifying spatial and temporal patterns of connectivity among populations is fundamental to marine ecology, evolutionary biology, and fisheries management. Yet, due to large population sizes and low genetic differentiation among populations, empirical quantification of population connectivity across a species’ entire range has not been achieved for an open-coast marine organism. Here, we leverage experimental transcriptomics to develop a genotyping-in-thousands by sequencing (GT-seq) panel to support assignment of recruits of the kelp forest gastropod, Kellet’s whelk (*Kelletia kelletii*), collected across the species’ biogeographic range. Over a three-year period, we identified high self-recruitment in the historical range (100%) and low self-recruitment in the expanded range (10.53 – 13.73%). Additionally, self-recruitment within the expanded range generally increased with recruit age, from 27.14% at 0.93 years to 43.40% at 1.93 years, indicating that the locally spawned individuals were more likely to survive to older ages than migrants from the historical range. Together, these results reveal limited self-recruitment in the expanded range and suggest that a post-settlement selective filter contributes to differential survival in a high gene flow marine system.

## Introduction

Identifying spatial and temporal patterns of connectivity among populations is fundamental to marine ecology and evolutionary biology, and critical for effective fisheries management and conservation (1,2). However, quantifying population connectivity and, by extension, delimiting boundaries among marine populations, can be challenging because dispersal often occurs during a pelagic larval stage where the larvae are translucent, miniscule (< 2 mm), and nearly impossible to track directly (3). This pelagic larval stage is ubiquitous; an estimated 80% of marine invertebrates and over 95% of all marine fishes have a pelagic larval stage as part of their life histories (4,5). Further, these larvae can be transported by ocean currents to populations that are hundreds of kilometers away from where they spawned (6,7), generating high levels of gene flow across diverse marine taxa (8–12). On the other hand, behavioral adaptations, homing mechanisms, and a complex interplay of biophysical processes can result in larvae exhibiting self-recruitment – that is, returning to the same population from where they were spawned (13–16). Consequently, the spatial scale and temporal variation in population connectivity is highly uncertain for the vast majority of coastal marine species (7).

Minuscule larvae and the large and variable spatial scales over which they disperse make traditional measures of population connectivity difficult to implement in most marine species (*i.e.,* mark-recapture). Additionally, large population sizes and low differentiation among marine populations impedes traditional *genetic* measures of population connectivity by reducing the power of parentage analyses and assignment tests, respectively (17,18). Empirical quantification of even one year of population connectivity across a species’ entire range has not been achieved for an open-coast marine organism (see 19,20 for singular examples in an estuarine and an amphidromous species using biochemistry of otoliths). Given the challenges associated with tracking larvae and the high variability in larval dispersal patterns, a novel and integrative approach is required to elucidate the degree to which marine populations are connected (21,22).

In high gene flow systems, signals of population differentiation can be swamped at neutral loci (23,24), and selection on standing genetic variation produces small peaks of differentiation (25), impeding population genetic analyses (26). However, larger signals have been identified at putatively adaptive loci under strong selective pressures (27–31). For example, signals of selection and differentiation in the coastal marine gastropod and commercial fisheries species, Kellet’s whelks (*Kelletia kelletii*), were revealed through an experimental transcriptomic approach that coupled common garden crosses and RNA sequencing (RNA-seq) to identify putatively adaptive loci (32,33). In contrast, previous analyses of this species based on neutral microsatellite loci failed to resolve population differentiation (11,12). Thus, using putatively adaptive loci instead of conventional neutral loci for conducting population assignment of recruits to natal sources offers an opportunity to directly measure population connectivity in coastal marine species with high gene flow (18).

Kellet’s whelk recently expanded its range distribution north, past a natural biogeographic break at Humqaq (Point Conception), California, USA, into an ecologically distinct, much colder bioregion (34–37). Genes associated with metabolism and cold tolerance were found to be upregulated in whelks from the expanded range (33). Population structure between the expanded and historical regions was identified at functional genes (*i.e.,* SNPs found across the transcriptome) and putatively adaptive genes (*i.e.,* SNPs found on DEGs) (33), despite the populations appearing to be panmictic at putatively neutral microsatellite loci (11,12).

Here, we leverage the population structure identified through differentially expressed genes to develop a genotyping-in-thousands by sequencing (GT-seq) panel to support assignment of Kellet’s whelk recruits to historical- vs. expanded-range spawning areas. Previous research has emphasized the importance of natural selection during larval dispersal and recruitment (7,38–41). To examine whether selection continues to shape the genetic makeup of cohorts after successful settlement (42), we also investigate patterns of survivorship as recruits matured. Altogether, we quantify regional connectivity and reveal the complex spatio-temporal eco-evolutionary dynamics of an open coast marine species.

## Methods

### Field collections

Using SCUBA, we collected adult (> 60mm shell length; 43) and age-0 to age-2 recruits (11.67 to 37.56mm; 44) Kellet’s whelks by hand from sub-tidal locations (approximately 15 m depth) across the species’ entire biogeographic range, from Isla San Roque, Baja California, Mexico to Monterey Bay, California, USA (CDFW SCP #8018 to C.W.). Collections occurred across three years during the summer months, from 2015 to 2017 (Figure 1, Table S1). DNA was extracted from whelk tissue using a Salting-out protocol (as described in 45), cleaned using the ZR-96 DNA Clean-Up Kit (Zymo Research, USA), and sequenced using the GT-seq panel described below.

**Figure 1.**
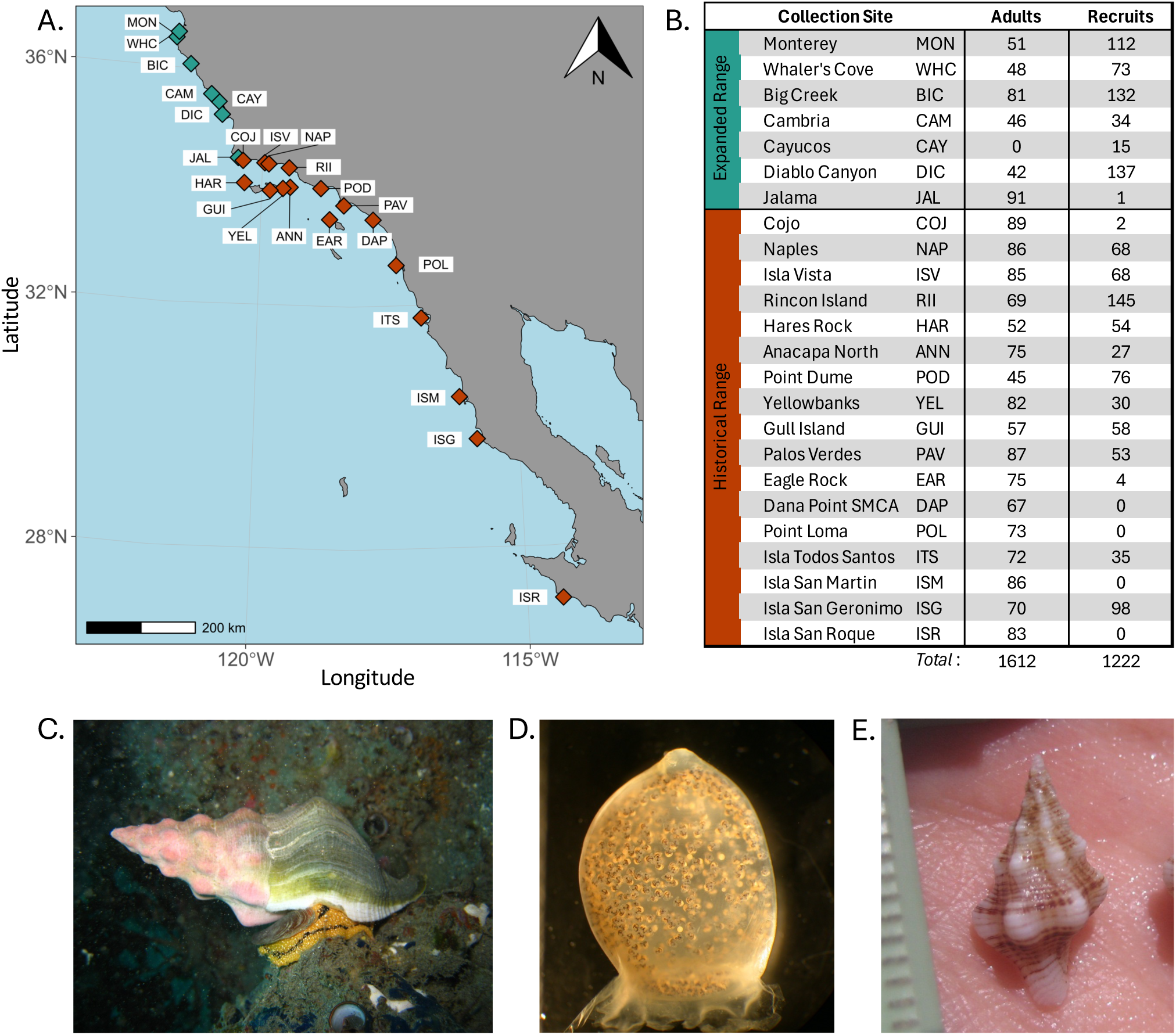
Sample collection sites and study species: A) Collection sites across Kellet’s whelks biogeographic range. B) Summary of samples for analysis at each collection site. Photographs of a Kellet’s whelk C) adult, D) egg capsule containing 500-1000 veliger larvae prior to hatching, and E) a recently-settled recruit.

### Sequencing and filtering

We developed a novel GT-seq panel (46) using single nucleotide polymorphisms (SNPs) found on differentially expressed genes (DEGs) between Kellet’s whelks’ expanded and historical range sites (MON and NAP, respectively; Figure 2) (33). Briefly, RNA reads were aligned to a *de novo* reference transcriptome (47) using bowtie2/2.4.2 (48), sorted using samtools/1.9 (49), and merged using stringtie2/2.1.1 (50). The count matrix was created using the featureCounts tool of subread/2.0.1 (51). DEGs were identified using the R package DESeq2/1.34 (52) using a minimum significance threshold of 0.05 after false discovery rate correction via the Benjamini–Hochberg method (53). SNPs were identified on DEG contigs using the GATK pipeline (54), following best practices for RNA-seq. We then selected 1,000 SNPs with the highest pairwise *F_ST_* values for GT-seq multiplex primer design by GTSeek (Twin Falls, ID).

**Figure 2.**
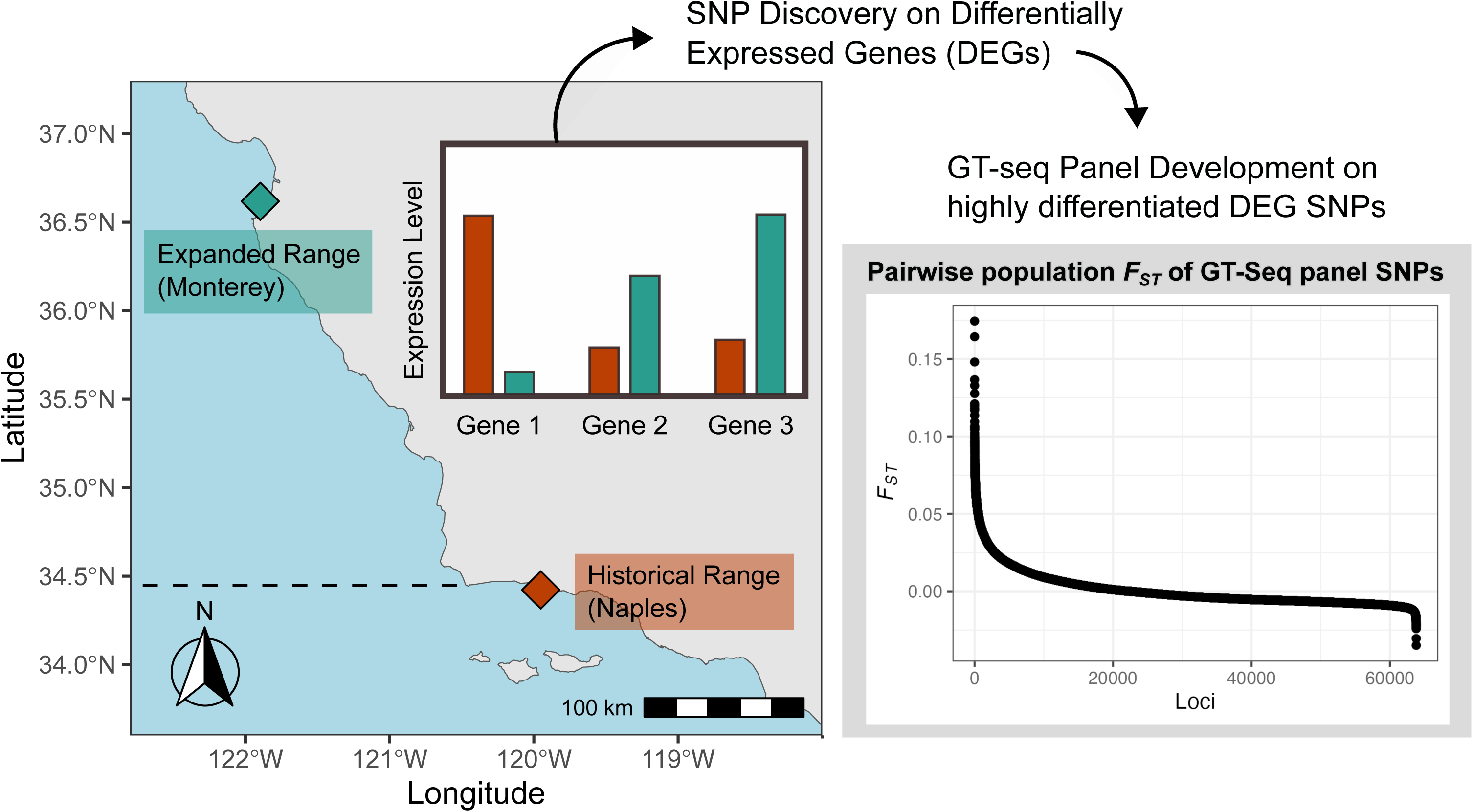
Study design and GT-seq panel development. Differentially expressed genes in Kellet’s whelk veligers from Monterey and Naples were identified in a common garden experiment (33); SNP variants were then discovered on these DEGs. The GT-seq panel was developed on highly differentiated SNPs. Pairwise population *F_ST_* of each GT-seq panel locus is shown in the gray panel (max *F_ST_ =* 0.1637).

### Genetic analyses of adults

Adult population structure is not expected to change substantially among consecutive years because Kellet’s whelks are long-lived and slow-growing (43). Consistent with this expectation, we observed minimal genetic differentiation between 2015 and 2016 at each site (median pairwise *F_ST_* between years = 0.00048, Figure S1). Accordingly, adult samples from both years were combined for all subsequent analyses. Prior to analysis, we first removed individuals with missing data in >20% of loci, then removed loci with missing data in >20% of individuals. To evaluate the genetic diversity across Kellet’s whelk’s entire biogeographic range, we calculated observed heterozygosity of each adult population using the *snpR* package (55). We also calculated mean relatedness at each collection site using Wang’s estimator (56) in the R package *related* (57). Results were visualized using *ggplot2* (58) in R (59).

### Population assignment of recruits

Population assignment tests were conducted using the infer_mixture() function in the R package *Rubias* (60) with 20,000 reps and 2,000 burn-in runs. All 1,612 adult individuals were used to create a reference for the expanded and historical regions (Figure 1, Table S1). Recruits from three consecutive collection years: 2015 (n = 363), 2016 (n = 482), and 2017 (n = 377) were used as the mixture (*i.e.,* samples to be assigned) (Figure 1, Table S1). We ran assignment tests using the default conditional Bayesian approach (method = “MCMC”) and the fully Bayesian approach (method = “BR”). Connectivity between regions was visualized using recruits with posterior means of group membership (PofZ) above 0.8 using the R package *ggalluvial* (61).

### Age-dependent self-recruitment

To understand patterns of survivorship as recruits matured, we quantified the age of each recruit in relation to its shell length using a growth function developed for the species (44), then investigated the relationship between age and self-recruitment, defined here as the proportion of recruits collected in a region that were also genetically assigned to that region. We calculated a moving average of self-recruitment proportion as a function of recruit age using a sliding window approach (window width = 0.6 years, step = 0.01 years). To assess the robustness of the assignment test, in addition to using samples with a posterior means of group membership (PofZ) value greater than 0.8, we repeated the analysis with larger sample sizes using recruits with PofZ values greater than 0.7.

To evaluate age-dependent survivorship to novel conditions (e.g., colder temperatures) in the expanded range, we calculated the proportion of self-recruitment as a function of age in four expanded populations with sufficient sample sizes: MON, WHC, BIC, and DIC. We used a PofZ cut-off of 0.7 for this analysis, as sample sizes were insufficient with a PofZ threshold of 0.8.

## Results

### Sample collection and classification

In total, our study analyzed 3,287 sequenced samples. After filtering out individuals with more than 20% of loci missing, 2,834 samples were retained for downstream analysis: 1,612 adult whelks collected from 23 sites in 2015 and 2016 (1,253 from the historical range and 359 from the expanded range) and 1,222 recruits collected from 20 sites in years 2015, 2016, and 2017 (Figure 1, Table S1).

### GT-seq Panel

Our GT-seq panel had 305 SNP loci. We removed 12 monomorphic loci and 31 loci missing in more than 20% of individuals, resulting in 262 SNPs for downstream analyses. Pairwise *F_ST_* values between collection sites for each locus ranged from −0.03 to 0.16 (mean = 0.00032, median = −0.0036, Figure 2). Pairwise *F_ST_* values between expanded and historical regions ranged from −0.0016 to 0.0073, with a mean of 0.00018 and a median of −0.00034.

### Adult populations

Mean relatedness within each collection site ranged from 0 to 0.22 (Figure 3, S3). Overall, relatedness was higher in the expanded range (mean = 0.0939) than in the historical range (mean = 0.0047). However, a few historical sites (COJ, NAP, ISV) closest to the expanded range also showed relatively higher relatedness (Figure 3). Observed heterozygosity at each collection site ranged from 0.1530 to 0.1676 (mean = 0.1608) and was similar between the expanded (mean = 0.1593) and the historical ranges (mean = 0.1613, Figure S4).

**Figure 3.**
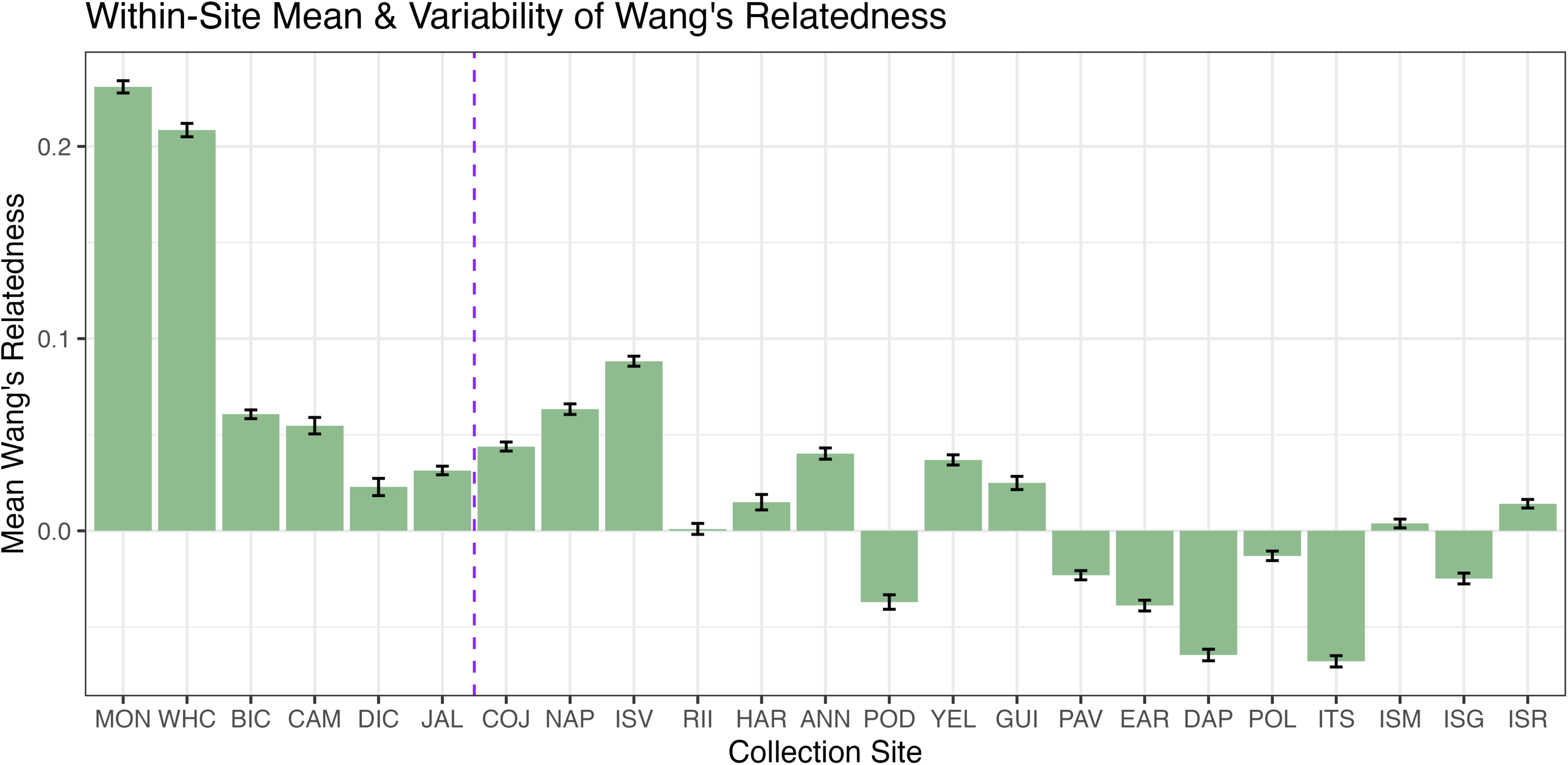
Relatedness in each adult population. Mean Wang’s relatedness in each population is plotted with standard error. Populations are ordered by latitude; the vertical dashed line separates the expanded (left) and historical region (right).

### Assignment of age-1 recruits

We used all 1,612 adult whelks to create a reference for population assignment. Of the 1,222 recruits collected across 3 years, 1,011 were classified as age-1 (Figure S2). We present assignment results for these age-1 individuals to understand processes that drive year-to-year variation in connectivity (Figure 4), although inclusion of age-0 and age-2 recruits did not alter general patterns of connectivity (Figure S5, S6). Of the age-1 recruits, 60.64% had posterior means of group membership (PofZ) > 0.8 (mean PofZ = 0.8163, Figure S7), resulting in the successful assignment of 608 age-1 recruits.

**Figure 4.**
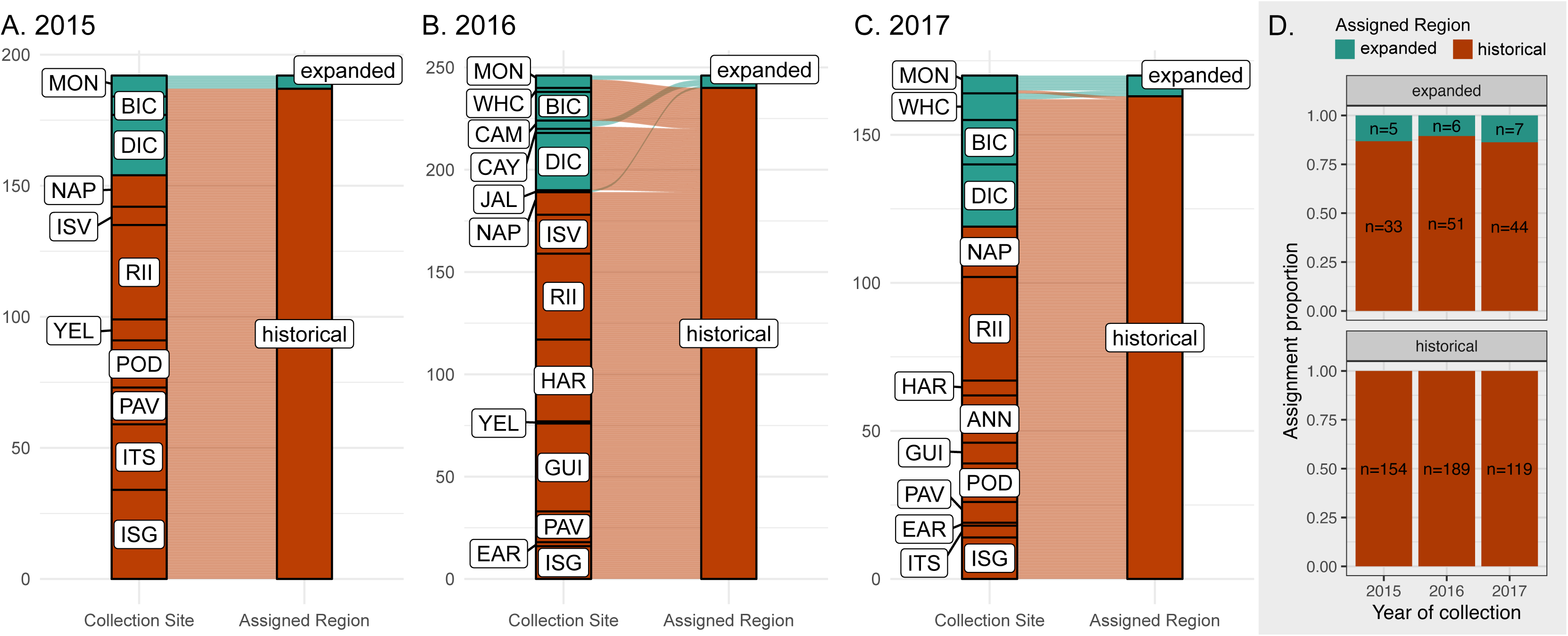
Genetic assignment of age-1 recruits. A-C shows genetic assignment results for age-1 recruits collected in 2015, 2016, and 2017 in alluvial plots. The left bar of each plot shows where each recruit was collected in the field, and the right shows the region (expanded or historical) where each recruit was genetically assigned. D) shows the proportion of recruit assignments to each region where the top panel shows recruits that were collected from the expanded region, and the bottom panel shows historical range recruits.

For age-1 recruits collected in the historical range, 100% (n = 462) recruited back to the historical range (*i.e.,* self-recruitment) (Figure 4). Assignment results did not significantly differ when using a conditional-Bayesian approach (Figure 4) or a fully-Bayesian approach (Figure S8). In contrast, the majority of recruits collected in the expanded range were not assigned back to the expanded range (12.33% self-recruitment, n = 146) (Figure 4). However, the assignment patterns were not consistent among all expanded range sites: every recruit collected from DIC and almost every recruit collected from BIC were assigned to the historical range, whereas as few as 17% of the recruits at MON (2017), the northernmost population in Kellet’s whelk’s expanded range, were assigned to the historical range (*i.e.,* 83% self-recruited).

Overall, self-recruitment in the expanded range was consistently low, regardless of collection year: 13.16% (n = 38) in 2015, 10.53% (n = 57) in 2016, and 13.73% (n = 51) in 2017, with 2016 showing the smallest proportion of self-recruits in the expanded range (Figure 4). The pattern of lower self-recruitment in 2016 was especially stark for recruits collected in MON: in 2015 and 2017, the majority of recruits were assigned to the expanded range; 62.5% (n = 8) and 83.33% (n = 6), respectively, whereas in 2016 self-recruitment declined to 33.33% (n = 8).

### Age-dependent self-recruitment

The vast majority of recruits collected in the historical range were assigned to the historical region (*i.e.,* self-recruitment), regardless of age (Figure 5). Conversely, self-recruitment within the expanded range increased with recruit age. To avoid obscuring range-wide patterns, Figure 5 excludes recruits collected in DIC, because DIC exhibited a broader age distribution than the other sites (Figure S9) and every DIC recruit, regardless of age, was assigned to the historical range (Figure 4). The proportion of self-recruitment in the expanded range increased from 27.14% at 0.93 years to 43.40% at 1.93 years (Figure 5). In other words, as a cohort of recruits from the expanded range aged and experienced mortality, a larger proportion of the surviving whelks in the cohort were assigned to the expanded range (Figure 5). These patterns were consistent across varying levels of assignment stringency (Figure S10). Additionally, we observed a latitudinal gradient of self-recruitment, where populations from more northern sites experienced higher levels of self-recruitment compared with populations further south in the expanded range (Figure S11).

**Figure 5.**
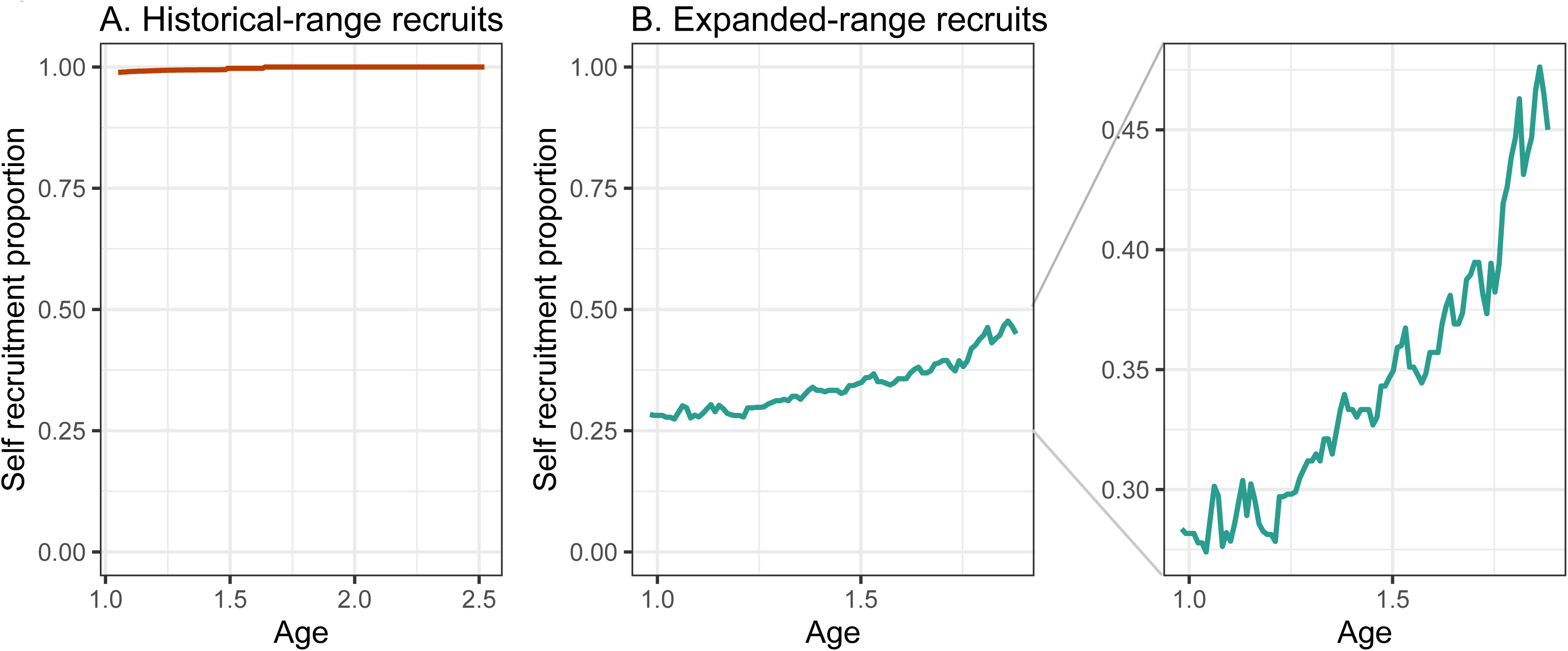
Relationship between age and self-recruitment (genetic assignment to the region of collection) for recruits (PofZ > 0.7) from A) the historical range and B) expanded range (excluding DIC). We exclude recruits collected in DIC, because DIC exhibited a broader age distribution than the other sites (Figure S9) and every DIC recruit, regardless of age, was assigned to the historical range (Figure 4). Self-recruitment was calculated using a sliding age window (window width = 0.6 years, step = 0.01 years), with values plotted at midpoints of each window.

## Discussion

Climate change is shifting the distributions of marine species and driving adaptation to novel environments (62–66), generating a natural experiment for studying the interplay between dispersal, gene flow, and genetic adaptation. Understanding these processes is also fundamental to marine science and critical for effective fisheries management and conservation (1,2,67–69). Our study quantified regional connectivity across the entire biogeographic range of a kelp forest gastropod and commercial fisheries species by leveraging a novel GT-seq panel based on differentially expressed genes (DEGs). This design allowed us to assign recruits collected from Kellet’s whelks’ entire range back to their natal region (expanded or historical range) across three consecutive years, revealing broad-scale patterns of connectivity across space and time (and their potential drivers).

Our intensive collection efforts of adults across 23 sites (n = 1,612) allowed us to understand how genetic diversity is distributed across the species’ range (Figure 1). Relatedness within populations was higher in the expanded range, suggestive of pulses of cohorts remaining together during dispersal (Figure 3). This result is consistent with previous hypotheses of long distance and episodic recruitment based on size-frequency distributions (37,70) and population genomics analysis at neutral loci (35) in Kellet’s whelk. The results also align with simulations of ocean currents in Kellet’s whelk’s range indicating larval dispersal to be intermittent and heterogeneous, causing population connectivity to be driven by packets of dispersing larvae between source and destination locations (71), and genetic analysis of fish species in Kellet’s whelk’s range finding high relatedness among cohorts of recruits (72).

The abundant center and central-marginal hypotheses both suggest that marginal populations are less genetically diverse and more likely to go extinct (73,74). This pattern may be further exacerbated by genetic drift and founder effects at range edges (75). Contrary to these predictions, we detected no reduction in genetic diversity in populations either at the leading edge of the expanded range or the trailing edge of the historical range (Figure S4).

Cross-recruitment from the historical range to the expanded range was the dominant pattern among the expanded range sites, indicating a large contribution of larvae from the historical range to the expanded range (Figure 4). Consistent dispersal and successful settlement of recruits past the historical range limit at Humqaq (Point Conception), California, USA (34,35,37) is suggestive of open populations and high gene flow in Kellet’s whelk. However, closer inspection reveals a latitudinal pattern in cross-recruitment: almost every recruit collected in expanded-range sites closest to the historical range (BIC and DIC) were assigned to the historical range, whereas cross-recruitment was much lower (thus self-recruitment much higher), in the northern most sites farthest from the historical range (MON and WHC; Figure 4, Figure S9).

Evaluation of assignments across three consecutive years with different oceanographic conditions enabled us to explore oceanographic drivers of population connectivity. In particular, interannual variation in oceanographic conditions may explain differences in recruitment patterns across years at the leading edge of Kellet’s whelk’s distribution in Monterey Bay, CA (MON). An analysis of age-1 recruits in this population reveals high levels of self-recruitment in 2015 and 2017, compared with high cross-recruitment from the historical range in 2016 (Figure 4), indicating enhanced long-distance dispersal that year. This cohort dispersed during late summer 2015 through early winter 2016, a period characterized by strong Pacific Decadal Oscillation (PDO) and El Niño–Southern Oscillation (ENSO) events (76). ENSO events alter current dynamics and are hypothesized to promote long-distance, northern larval transport in California (37,70,71,77,78), but see (79). This pattern is consistent with previous population genetic studies in this species (35), and a length–age model for Kellet’s whelk indicating that the first individuals found in MON and BIC spawned between 1969 and 1974, coincident with the 1972–73 ENSO event (44). Combined, these findings suggest that ENSO may have enhanced long-distance, northern dispersal to MON in 2016.

As recruits collected in the expanded range aged, a larger proportion of the individuals in the cohort were assigned to the expanded range (Figure 5). This shift toward self-recruitment with time indicates that the locally spawned (self-recruited) individuals were more likely to survive to older ages than migrants from the historical range. This pattern reveals a post-settlement selective filter in the expanded region. We hypothesize that this pattern is due to viability selection in favor of locally born individuals that may be genetically adapted to novel conditions (e.g., colder temperatures) in the expanded range (33).

Previous research has emphasized the importance of natural selection during larval dispersal and recruitment (7,38–41). For the vast majority of coastal marine fish and invertebrate species, ecological theory posits this short period of larval settlement as a critical bottleneck, where natural selection causes >99% mortality due to predation, starvation, and habitat factors (4,80,81). Following this bottleneck, the surviving recruits have developed sufficient size and defenses to largely avoid mortality and grow up into juveniles and then reproductive adults. Consequently, this pre-settlement bottleneck is thought to be the major determinant of the species’ adult population genetic structure (7,38,39,41). Comparatively few studies have extended this hypothesis to post-settlement stages (82–84) – such that juvenile mortality further changes population genetic structure (42,85,86). Among the few studies to explicitly explore this topic, (86) and (42) found post-settlement mortality during the juvenile stage to significantly alter the genetic structure of the subsequent adult populations (in a fish from the Mediterranean Sea and an intertidal crab from the northeastern Pacific, respectively). Here we build upon these studies and explore post-settlement mortality in an open-coast sub-tidal species that exhibits high levels of gene flow. We provide additional support for the hypothesis that differential survival of settlers can generate a pattern of *realized* population connectivity of established juveniles (older recruits) that is structurally different from the pattern of *potential* connectivity estimated from newly settled individuals (younger recruits) (Figure 5).

Our targeted sequencing panel based on DEGs allowed for the first empirical quantification of broad-scale connectivity across an open-coast marine species’ range — helping to crack open the population connectivity “black box” (*sensu* (87)). Although our approach enabled robust assignment at the regional scale, resolving connectivity at the population level may require genome-wide marker sets, loci with more discriminatory power, and larger reference panels (17,88,89). Recent developments in population assignment tests that leverage data from across the genome (90,91) may offer increased power to identify connectivity at a finer scale in high gene flow species in future studies. Additionally, our GT-seq reference panel had higher confidence when assigning recruits to the historical range (Figure S5), which may be partially driven by differences in sample sizes. Consequently, recruits from the expanded range had lower average PofZ values and were more frequently filtered out (Figure S5).

Our findings provide new insights into a fundamental question in marine ecology: Are marine populations open or closed? Dispersal, recruitment, and gene flow from the historical range to the expanded range is very high (Figure 4), suggestive of open populations in Kellet’s whelks. However, following settlement, there is a strong selective filter in the expanded range, such that not all individuals who successfully disperse to and settle in the expanded range survive to adulthood (Figure 5). Thus, while populations may appear demographically open when analyzed using conventional population genetic approach (11,12), realized genetic connectivity is markedly more closed due to differential survival after settlement. This study reveals how population differentiation can occur in high gene flow systems by identifying a post-settlement selective filter. Thus, range expansion in Kellet’s whelk is shaped not solely by dispersal dynamics, but also by viability selection after arrival. These findings emphasize the importance of regional genetic adaptation in mediating gene flow during range expansion, and challenge the long-standing assumption in marine ecology that heavy, selective mortality during dispersal and initial settlement alone determines the genetic structure of coastal marine populations (38,41,92).

## Notes

https://www.bco-dmo.org/dataset/995189

### Competing Interest Statement

The authors have declared no competing interest.

### Funding

- U.S. National Science Foundation: OCE-1924537, OCE-1924505, OCE-1924604

