## Supplemental Information for "Regional connectivity and viability selection in a range-expanding marine species"

Table S1

Figures S1 to S11

**SI Tables**

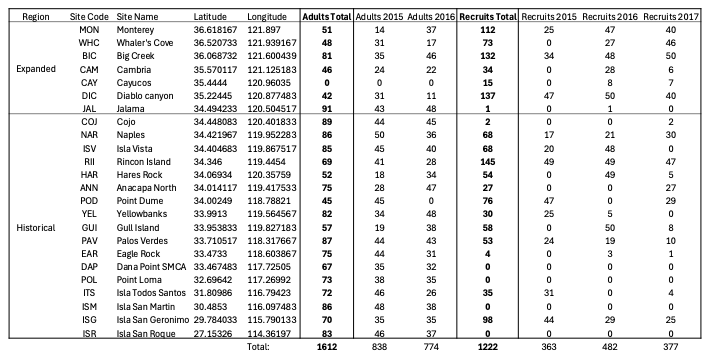
**Table S1**. Sample sites and expanded collection summary, listing each collection site’s region (expanded or historical), abbreviation (Site Code), name, latitude, longitude, and number of adult and recruit collected in each year.

SI Figures

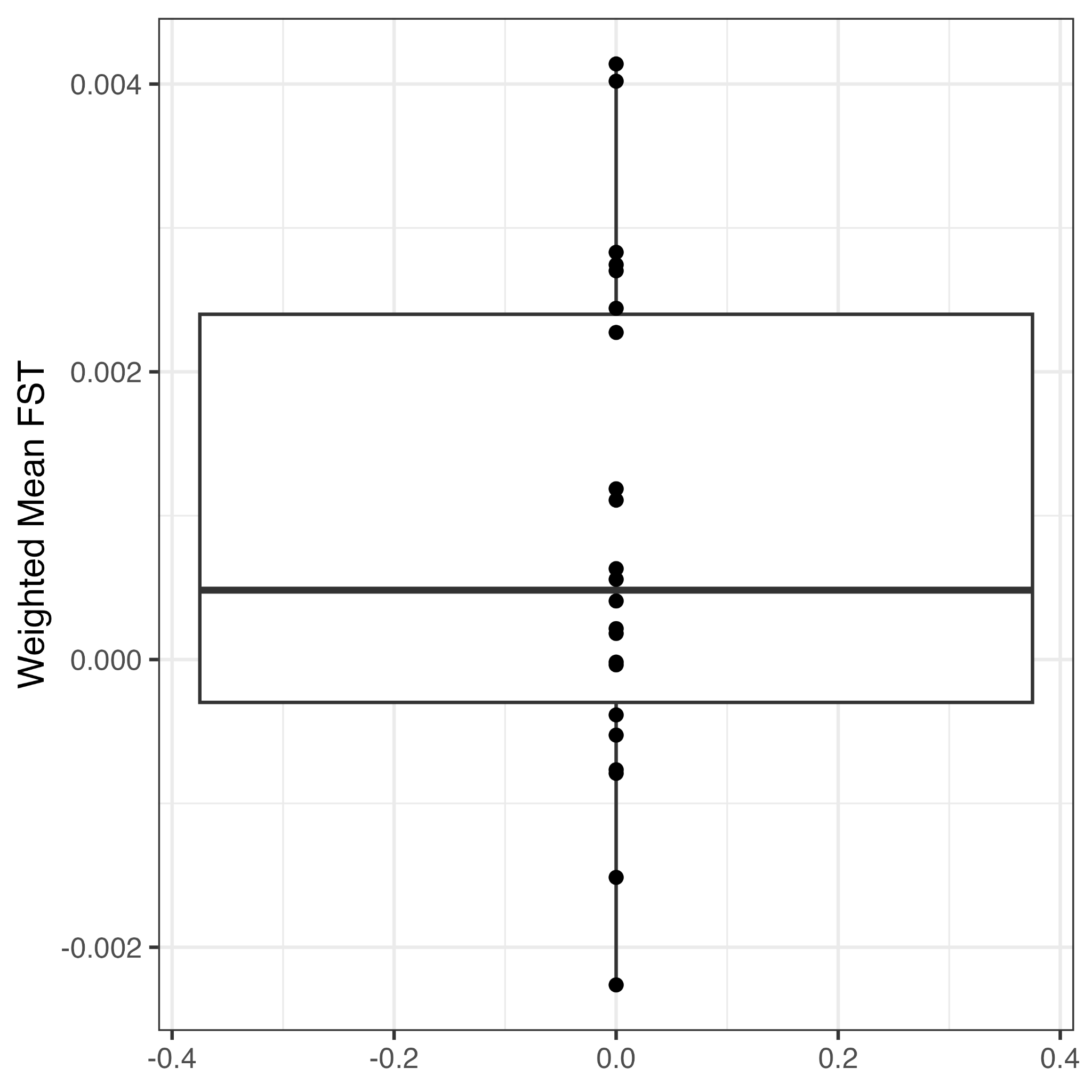

**Figure S1.** Pairwise weighted mean *F_ST_* between 2015 and 2016 adult collections at each site. *F_ST_* values range from -0.0023 to 0.0041, with a median of 0.00048 and a mean of 0.00087.

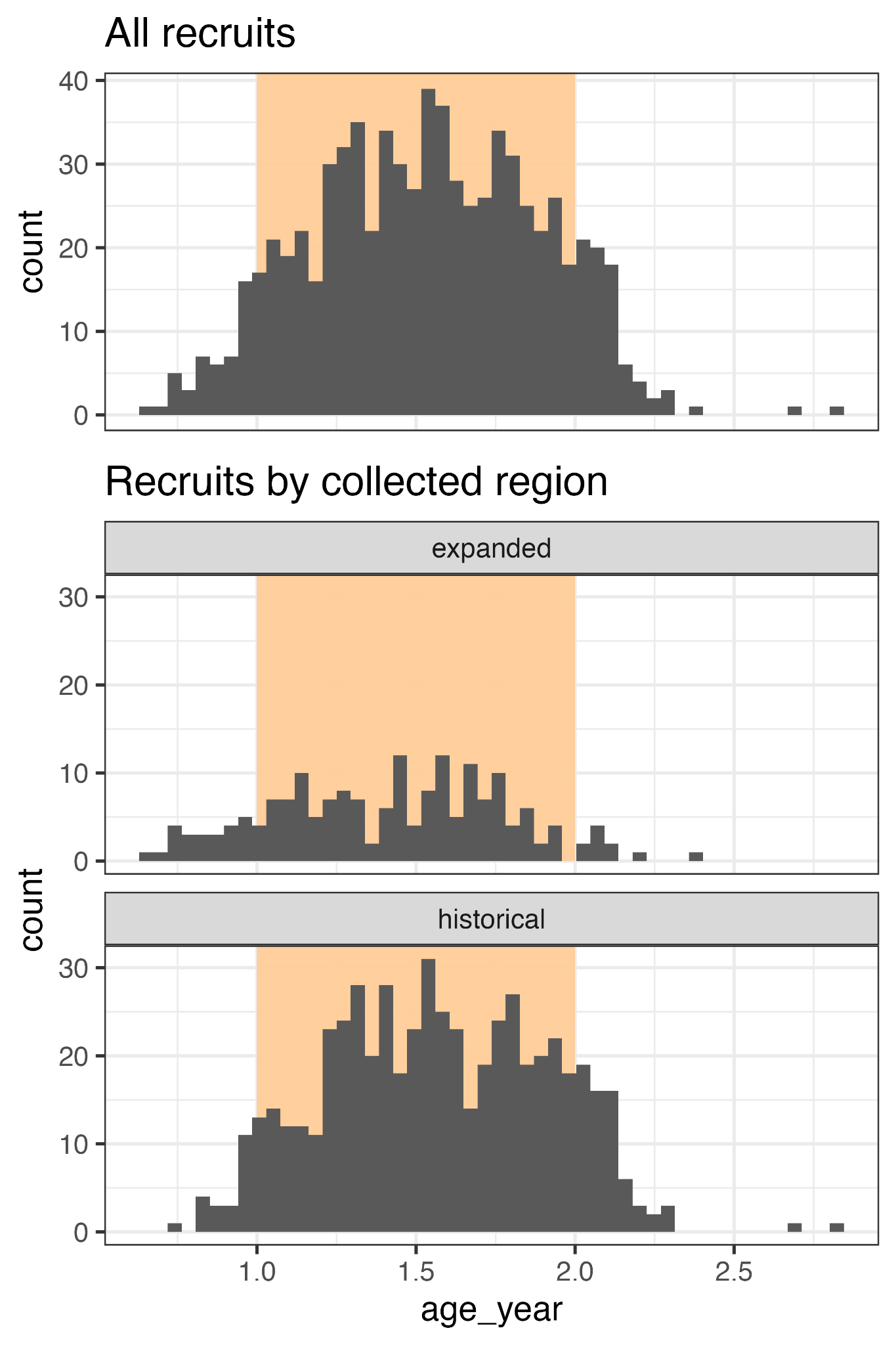

**Figure S2**. Histogram of recruit ages. Light orange backgrounds indicate age-1 recruits.

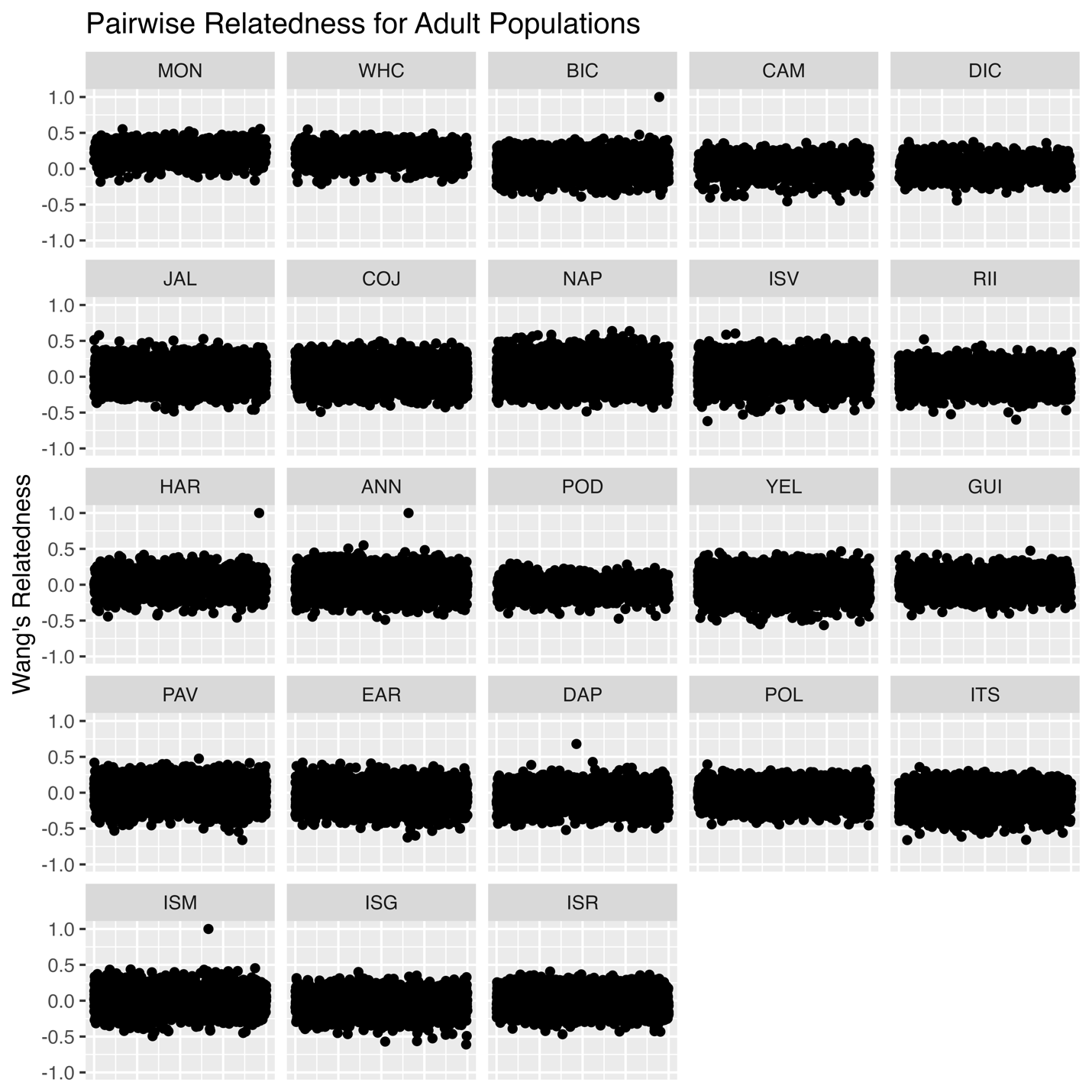

**Figure S3**. Pairwise relatedness for adult samples. Four samples (one each from BIC, HAR, ANN, and ISM) were collected across years from the same site and represent re-captured individuals.

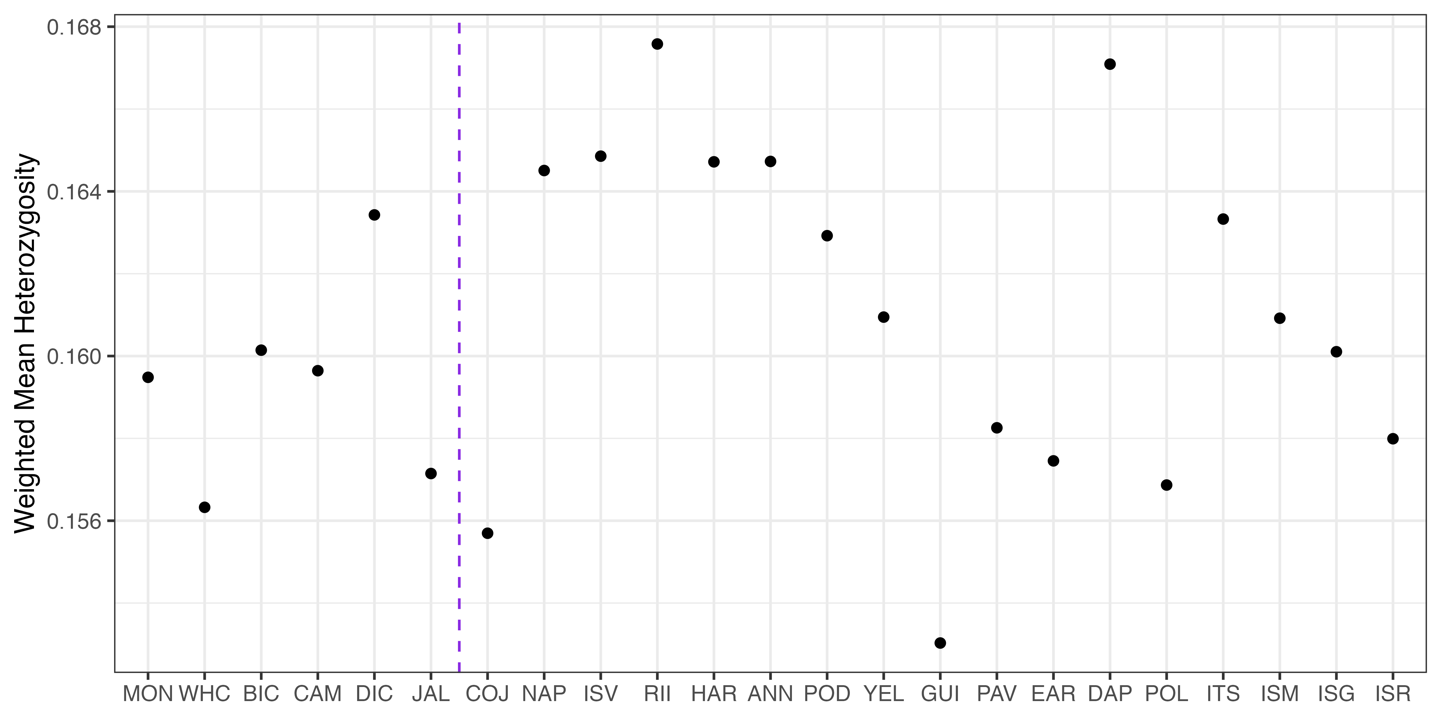

**Figure S4**. Observed heterozygosity of each adult population. Populations are ordered by latitude, the purple dashed line shows the expanded (left) and historical region (right).

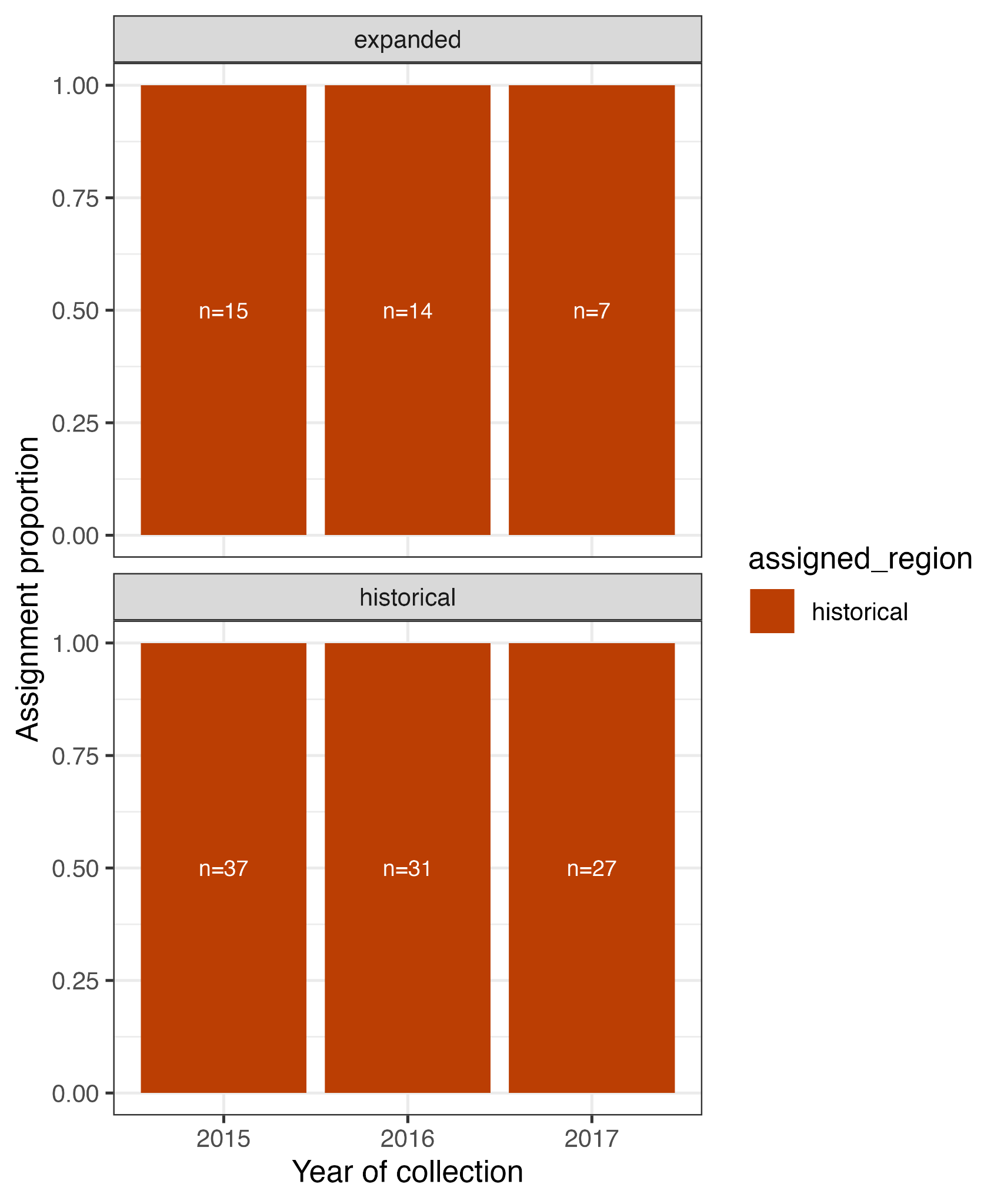

**Figure S5.** Recruit assignments for age-0 and age-2 recruits using a fully conditional approach in Rubias (PofZ > 0.8). The top panel shows recruits that were collected from the expanded region and the proportion that was assigned to expanded or historical range (shown in colors), the bottom panel shows historical range recruits.

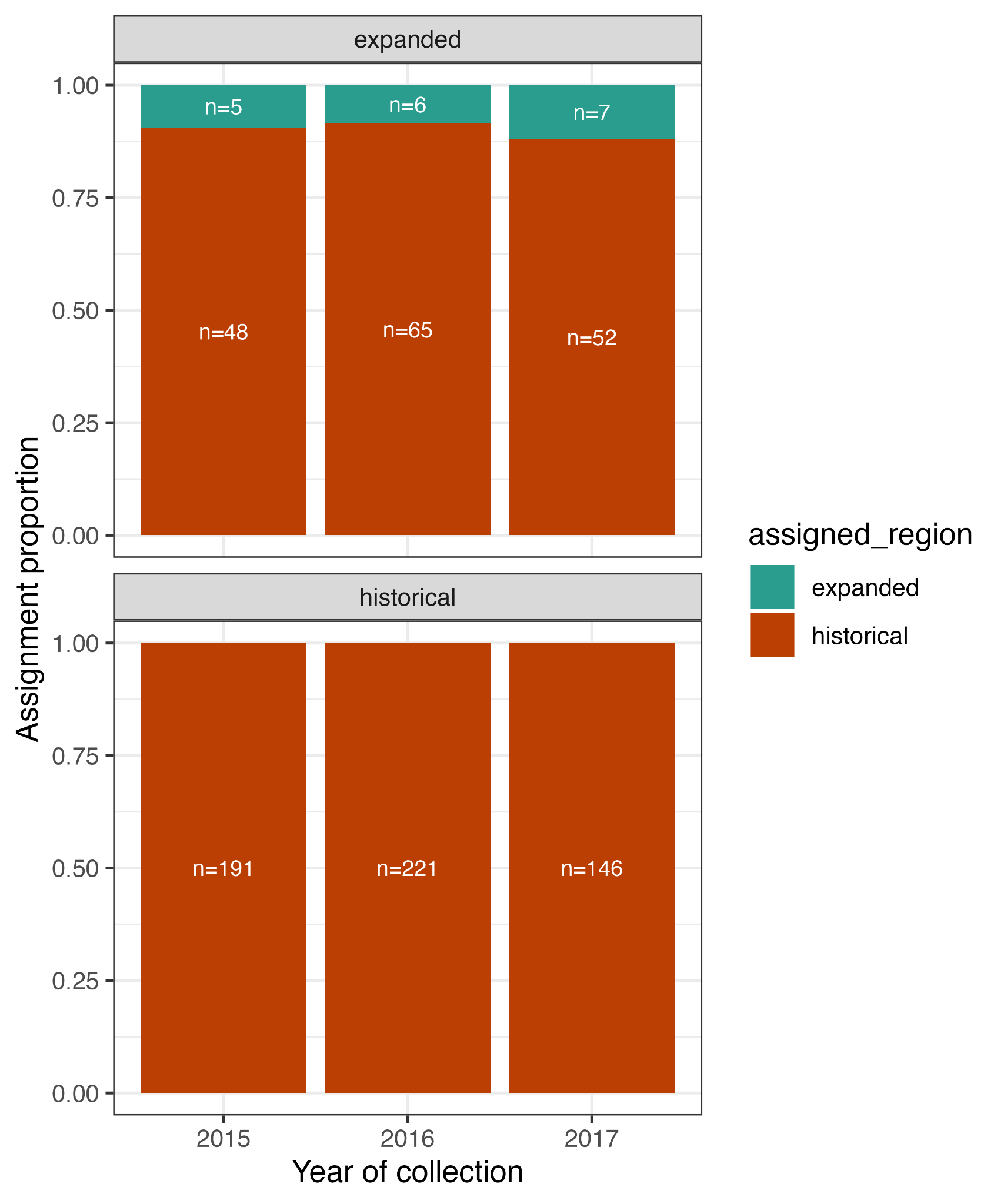

**Figure S6.** Recruit assignments for all individuals, including age-0 and age-2 recruits using a conditional Bayesian approach in Rubias. The top panel shows recruits that were collected from the expanded region and the proportion that was assigned to expanded or historical range (shown in colors), the bottom panel shows historical range recruits.

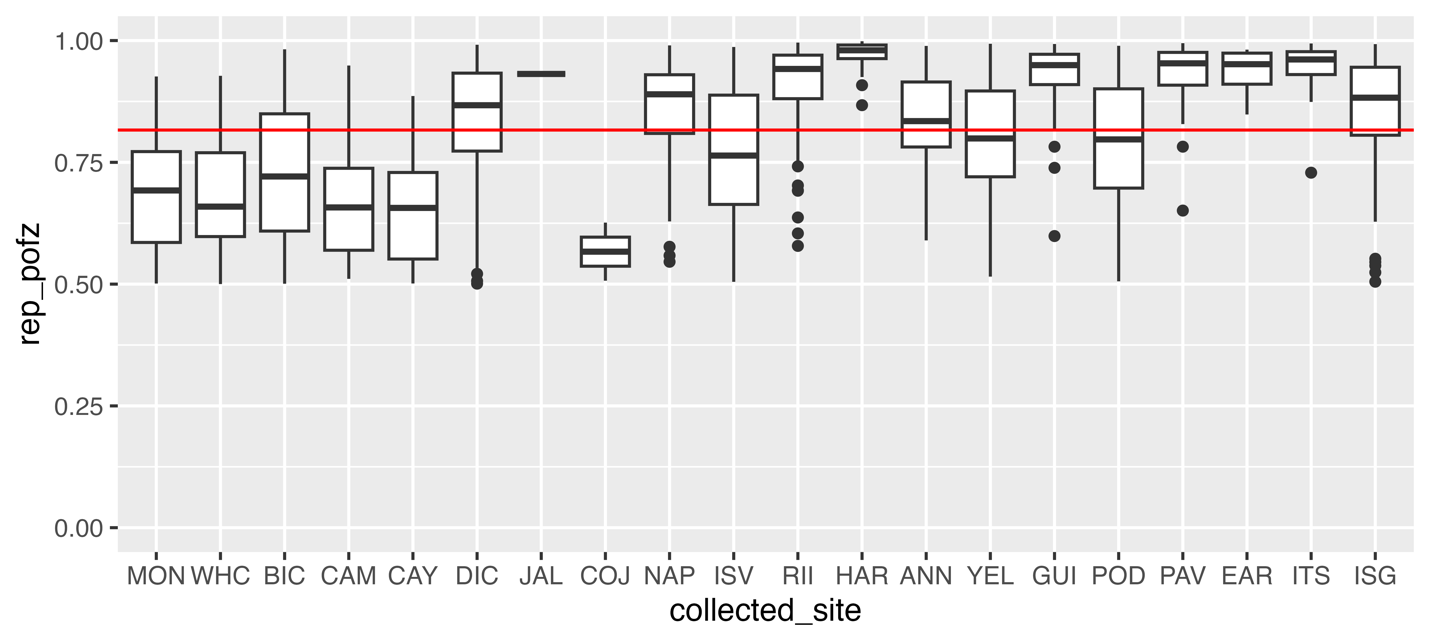

**Figure S7.** Mean posterior means of group membership (expanded or historical range) for all recruits at each collection site. The horizontal red line represents a PofZ cut-off at 0.8.

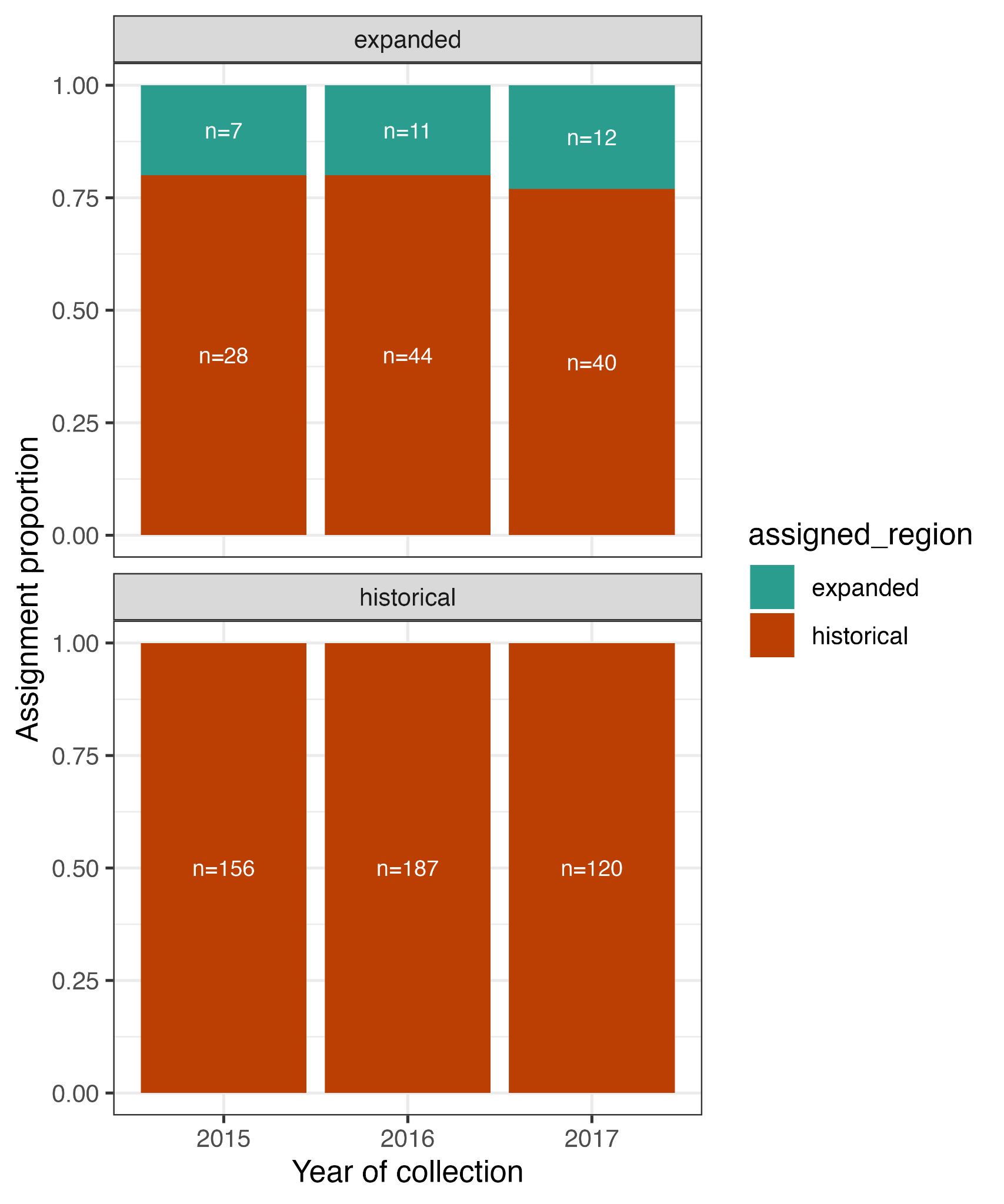

**Figure S8.** Recruit assignments using a fully Bayesian approach in rubias (method=“BR”). The top panel shows recruits that were collected from the expanded region and the proportion that was assigned to expanded or historical range (shown in colors), the bottom panel shows historical range recruits.

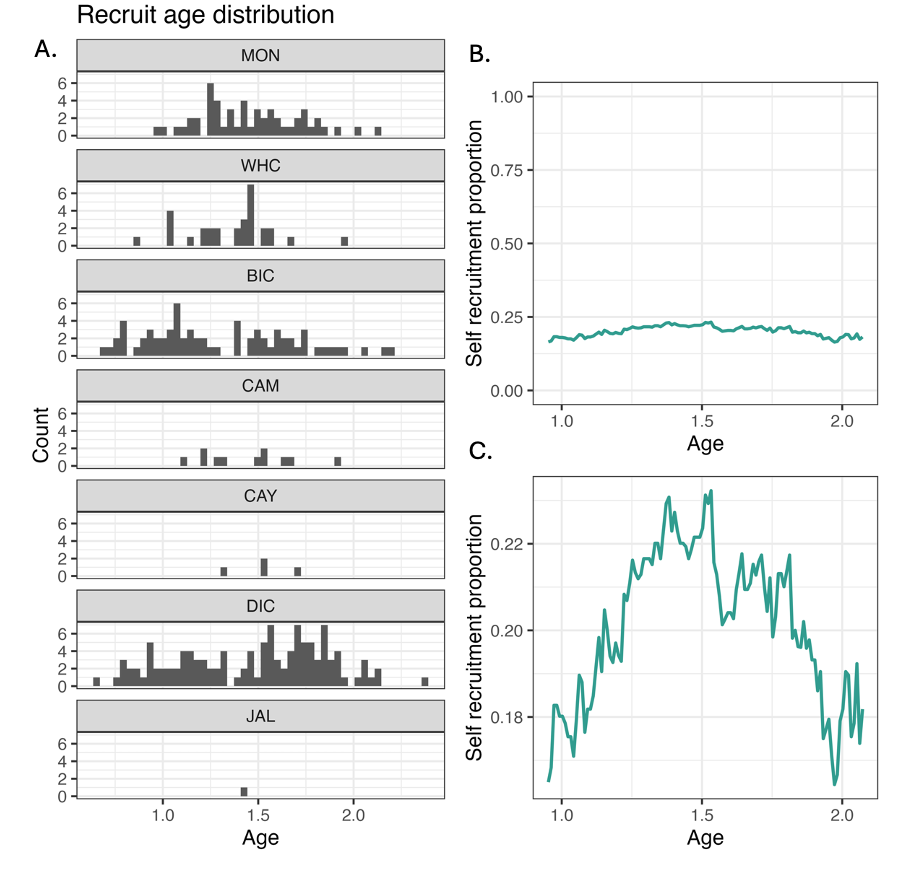

**Figure S9**. Effect of DIC on self-recruitment calculations. A) Recruit age distribution of each expanded population (PofZ > 0.7). B) Relationship between age and self-recruitment in expanded range recruits, inclusive of DIC. C) Zoomed-in view of relationship between age and self-recruitment.

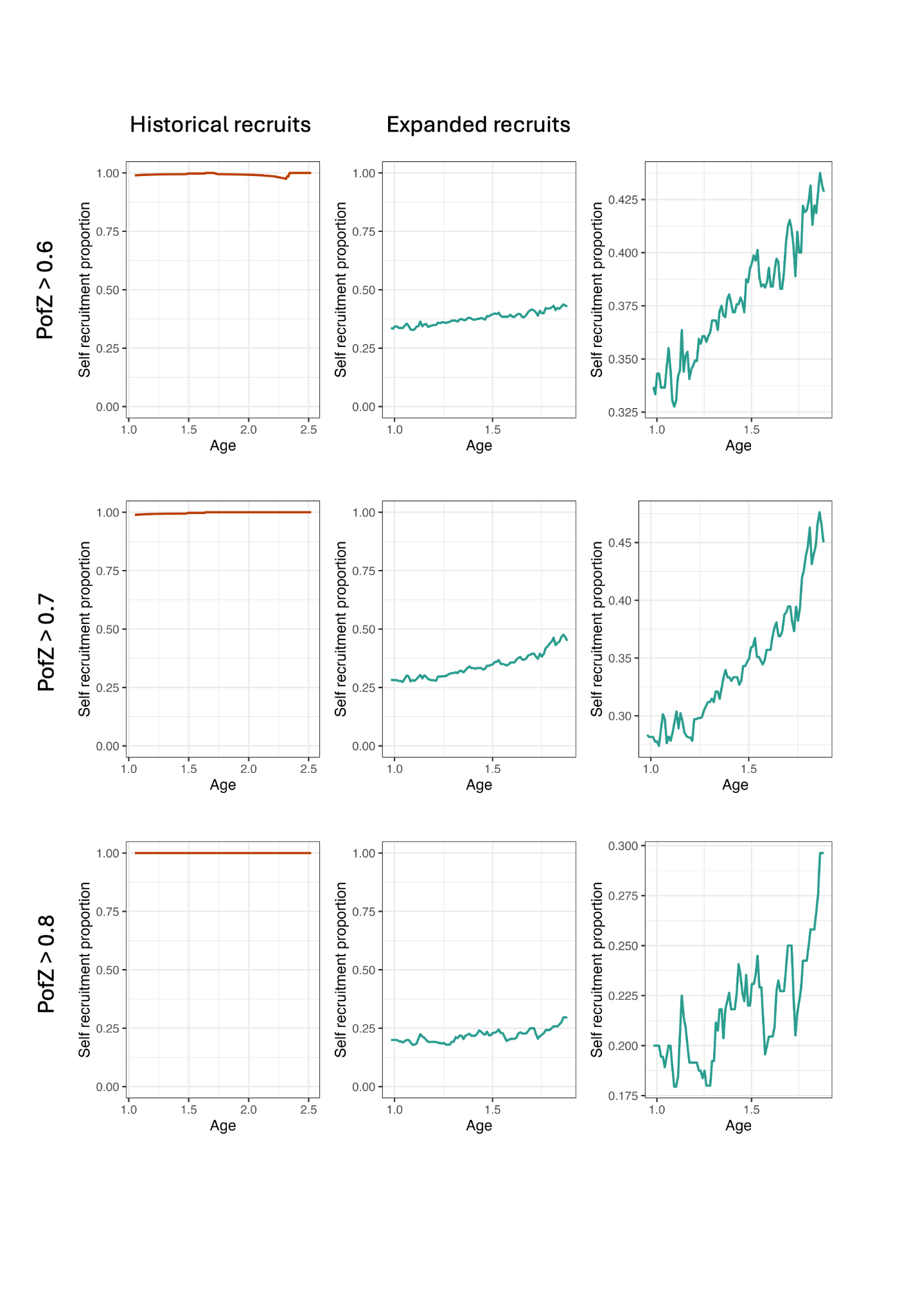

**Figure S10**. Relationship between age and self-recruitment in historical vs. expanded range recruits (all expanded populations except DIC). Self-recruitment was calculated using a sliding age window (window width = 0.6 years, step = 0.01 years), with values plotted at midpoints of each window. Proportion of self-recruitment was calculated for recruits with genetic assignment confidence (Rubias PofZ) values greater than 0.6, 0.7 and 0.8.

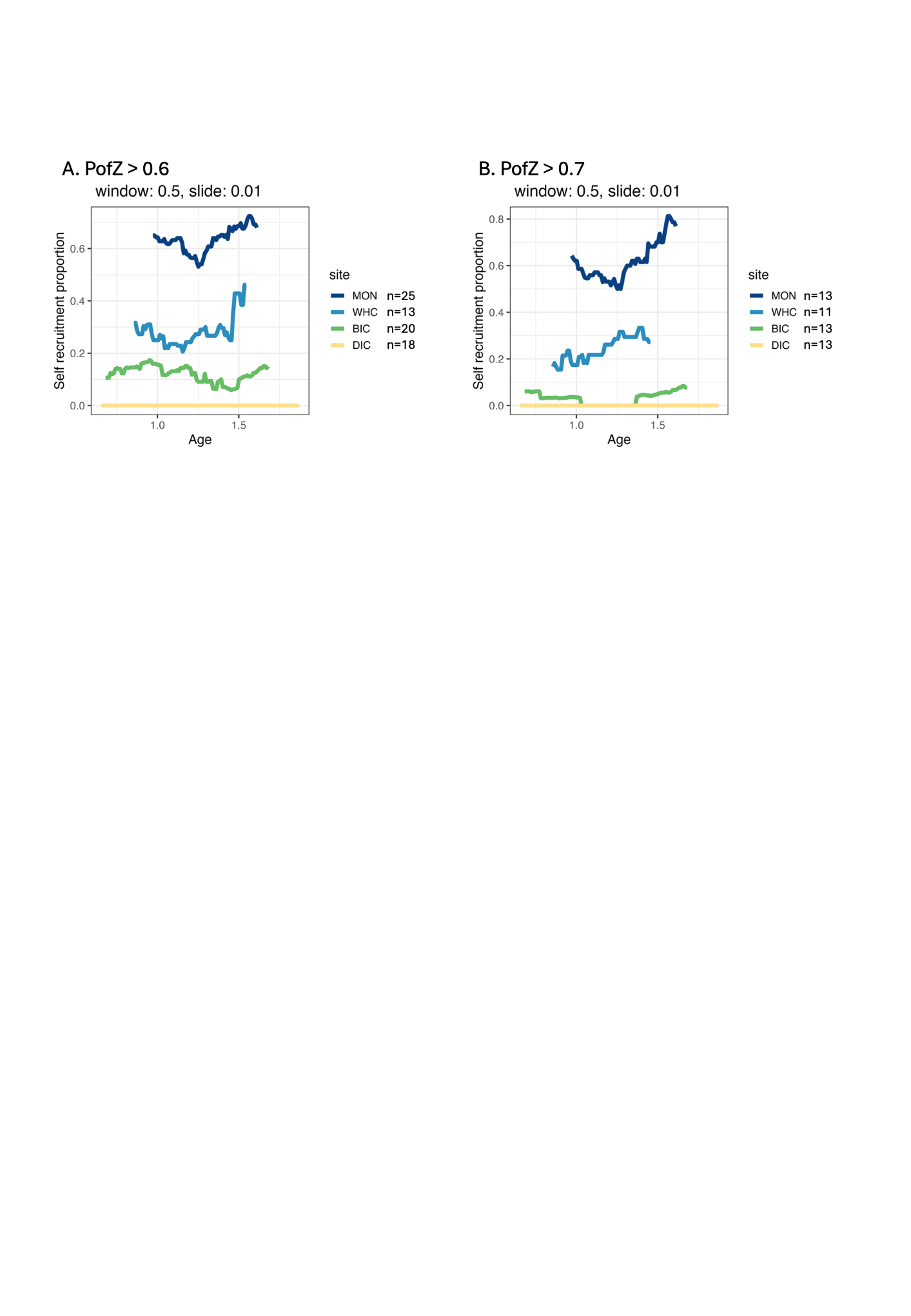

**Figure S11.** Relationship between age and self-recruitment in four expanded populations (MON, WHC, BIC, and DIC). Proportion of self-recruitment follows a latitudinal gradient. It is highest in MON, the northernmost population. Self-recruitment was calculated using a sliding age window (window width = 0.5 years, step = 0.01 years), with values plotted at midpoints of each window. The number of individuals present in each window is shown in the legends. Proportion of self-recruitment was calculated for recruits with genetic assignment confidence (Rubias PofZ) values greater than 0.6 (A) and 0.7 (B). There were insufficient sample sizes with a PofZ cut-off of 0.8.
